# TMEM100 attenuates NF-κB activation via disrupting the PRDX1-GNAI2 complex to alleviate acute lung injury

**DOI:** 10.1101/2025.10.24.684325

**Authors:** Cheng Qian, Linxin Pan, Chuncan Si, Youwen Du, Jingying Wang, Yue He, Bingjie Liu, Sa Xiao, Yufeng Zhu, Fengsong Wang, Kezhen Wang

**Author notes:** Corresponding author: Fengsong Wang,; Kezhen Wang. These authors contributed equally to this work (Cheng Qian, Linxin Pan).

## Abstract

Acute lung injury (ALI) is a severe and diffuse inflammatory disorder of interstitial lung. Emerging evidence suggests that TMEM100 is closely associated with lung development and function. However, its role in ALI remains unclear. In this study, we observed a significant downregulation of TMEM100 expression in both mouse lung tissues with ALI and lipopolysaccharide (LPS)-induced pulmonary vascular endothelial cells (PVECs). Overexpression of TMEM100 markedly attenuates LPS-induced lung injury and inflammation, while also restoring the imbalance between proliferation and apoptosis in PVECs. Mechanistically, TMEM100 interacts with both PRDX1 and GNAI2, disrupting the PRDX1-GNAI2 complex and thereby inhibiting LPS-induced NF-κB activation, which contributes to its anti-inflammatory effects. These findings highlight the protective role of TMEM100 in endotoxin-induced ALI and provide a theoretical basis for understanding its biological functions and potential applications in ALI gene therapy.

## Introduction

Acute lung injury (ALI) and its more severe form, acute respiratory distress syndrome (ARDS), are life-threatening inflammatory lung disorders characterized by high morbidity and mortality in critically ill patients [1]. Key pathophysiological features include elevated extravascular lung water, decreased lung volume and compliance, an impaired ventilation-perfusion ratio, and progressive hypoxemic respiratory failure. The pathogenesis of ALI involves the loss of alveolar–capillary membrane (ACM) integrity and the release of pro-inflammatory mediators [2]. As essential components of the ACM, pulmonary vascular endothelial cells (PVECs) are not only the first cells to be injured but also active participants in the inflammatory response, playing a pivotal role in the progression of ALI and related pulmonary circulatory diseases [3, 4].

Bacterial infections, particularly sepsis and severe pneumonia, are major triggers of ALI. During severe infections, lipopolysaccharide (LPS)-a key component of Gram-negative bacterial cell walls—is released into the bloodstream and recognized by cell membrane receptors, initiating targeted cell damage [5]. Structurally, LPS consists of lipid A, core polysaccharides, and O-antigen, with the latter determining its serotype and biological activity [6]. LPS-induced ALI involves multiple mechanisms, including inflammation, oxidative stress, and apoptosis, among which inflammation has been the most extensively studied [7]. Upon entering the circulation, LPS binds to lipopolysaccharides binding protein (LBP) and CD14, forming the LPS-LBP-CD14 complex. This complex, aided by myeloid differentiation protein-2 (MD-2), activates Toll like receptors 4 (TLR4), triggering intracellular signaling through myeloid differentiation primary response protein 88 (MyD88) and interleukin-1 receptor-associated kinase (IRAK) [8]. Subsequent downstream activation of the nuclear factor kappa-B (NF-κB) and mitogen activated protein kinase (MAPK) pathways via the adaptor protein tumor necrosis factor receptor-associated factor 6 (TRAF6) amplifies the inflammatory response, leading to pro-inflammatory cytokine release and the development of ALI [9, 10].

Transmembrane protein 100 (TMEM100), located on chromosome 17q32, is a conserved protein with two predicted transmembrane domains (53-75 and 85-107) [11]. It is expressed in multiple tissues, including the brain, heart, kidneys, prostate, lungs and liver, with the highest expression in the lungs. TMEM100 has been implicated in diverse biological processes, such as angiogenesis, neurodevelopment, pain transmission, and tumor suppression [12–16]. Database analyses (e.g., HPA, FANTOM5 and GEO) reveal that TMEM100 is highly expressed in normal lung tissues, but significantly reduced under pathological conditions such as mechanical ventilation-induced injury, inflammation, and cancer. Notably, TMEM100 exhibits anti-cancer effects in non-small cell lung cancer (NSCLC) and may serve as a prognostic marker [17, 18]. Recent studies have identified TMEM100 as an endothelial-enriched gene critical for pulmonary vascular repair and lung function maintenance [19, 20]. Our previous research also demonstrated its anti-inflammatory role in liver injury models [21], promoting further investigation into its function in ALI.

This study investigates the role of TMEM100 in the pathogenesis of ALI. Our findings demonstrated that TMEM100 expression was downregulated in mouse models of ALI and LPS-induced PVECs. Furthermore, overexpression of TMEM100 markedly alleviated LPS-induced lung injury, inflammation and PVEC dysfunction by inhibiting NF-κB activation. Mechanistically, TMEM100 was found to disrupt the PRDX1-GNAI2 complex, thereby suppressing the downstream NF-κB signaling pathway. Collectively, these results underscore the protective role of TMEM100 in ALI and highlight its potential as a therapeutic target.

## Materials and Methods

### Reagents and Antibodies

Fetal Bovine Serum (FBS) (FB25015) was purchased from Clark. LPS (HY-D1056) was purchased from MedChemExpress. The Lipo8000™ Transfection Reagent (C0533), RNAeasy™ Plus Animal RNA Isolation Kit (R0032), Cell lysis buffer for Western and IP (P0013), Protease Inhibitor Cocktail (P1005), Immunoprecipitation Kit with Protein A/G Magnetic Beads (P2179S), Primary Antibody Dilution Buffer (P0023A), and BeyoClick™ EdU-555 Cell Proliferation Kit (C0075S) and DAPI (C1006) were obtained from Beyotime Biotechnology. Antibodies against TMEM100 (25581-1-AP), TNF-α (17590-1-AP), IL-6 (21865-1-AP), GNAI2 (11136-1-AP), PRDX1 (15816-1-AP), and β-actin (66009-1-Ig) were sourced from Proteintech. The ECL Chemiluminescent Kit (P2200) was supplied by NCM Biotech. Evo M-MLV RT Premix (AG11706) and SYBR Green® Premix Pro HS qPCR Kit (AG11701) were obtained from AG. Mouse TNF-α (MU30030) and IL-6 ELISA Kit (MU30044) were purchased from Bio-swamp. The PE Annexin V Apoptosis Detection Kit I (559763) were acquired from BD Pharmingen.

### Animals and LPS-induced ALI Model

C57BL/6J mice (weighting 18-20 g) were obtained from Yongkang Qingyuan Farm (Dingyuan, China) and maintained under specific-pathogen-free conditions. All animal experiments were conducted in accordance with protocols approved by the Animal Care and Use Committee of Anhui Medical University (Approval No. LLSC20201078).

To establish the ALI model, mice received an intratracheal instillation of LPS at a dose of 10 mg/kg. For gain-of-function, AAV9 carrying TMEM100 (AAV9-TMEM100) or the corresponding control (AAV9-NC) was administered injection (1×10¹¹ viral genomes per mouse) three weeks prior to LPS challenge. All mice were euthanized 24 h after LPS administration for subsequent tissue collection and analysis.

### Cell Culture and Transfection

PVECs were acquired from Shanghai Yaji Biotechnology Co., Ltd. and cultured in Dulbecco’s Modified Eagle’s Medium (DMEM) supplemented with 10% fetal bovine serum (FBS) and 1% penicillin/streptomycin. Cells were maintained at 37°C in a humidified incubator with 5% COC.

For experimental treatments, HULEC-5A cells were seeded in 6-well plates and divided into four groups: Control, LPS (1 μg/mL), LPS + Vector (empty pcDNA3.1), and LPS + TMEM100 (pcDNA3.1-FLAG-TMEM100). Expression vectors were transfected using a liposome-based transfection reagent. After 6 h, cells were stimulated with LPS. Cells were harvested at 24 h post-transfection for subsequent Western blot or RT-qPCR analysis.

### Immunohistochemistry (IHC)

Lung tissue sections were deparaffinized in xylene and rehydrated through a graded alcohol series. Endogenous peroxidase activity was quenched by incubation with 3% hydrogen peroxide (H_2_O_2_), followed by blocking of nonspecific binding sites using pre-immune rabbit serum. Sections were then incubated overnight at 4°C in a humidified chamber with the following primary antibodies diluted in antibody diluent: anti-TMEM100 (1:50), anti-TNF-α (1:50), and anti-IL-6 (1:50). After three washes with phosphate-buffered saline (PBS), the sections were incubated with an HRP-conjugated secondary antibody for 30 min at room temperature. Antigen-antibody complexes were visualized using 3,3’-diaminobenzidine (DAB) substrate for approximately 3 min. For negative controls, the primary antibody was replaced with PBS. Finally, all sections were counterstained with hematoxylin, and digital images were acquired using a Tissue FAXS Plus S quantitative analysis system (Tissue Gnostics).

### Real-time Quantitative PCR (RT-qPCR)

Total RNA was extracted from lung tissues or cultured cells using TRIzol reagent. cDNA synthesis was synthesized from the extracted RNA using the Evo M-MLVRT Master Mix, according to the manufacturer’s instructions. Quantitative PCR (qPCR) was subsequently performed using the SYBR^®^ Green Premix Pro Taq HS qPCR Kit on an ABI Quant Studio™ 6 Flex Real-Time PCR system. Equal amounts of cDNA were applied for each reaction. The specific sequences of all oligonucleotide primers used in this study are listed in **Table S1**. Gene expression levels were normalized to the endogenous control β-actin and calculated using the comparative 2^−ΔΔCT^ method.

### Western Blot Analysis

Total protein was extracted from lung tissues or cultured cells using RIPA lysis buffer containing a protease inhibitor cocktail. Protein concentrations were determined, and equal amounts (30 μg per lane) were separated by 12% SDS-PAGE and subsequently transferred to Immuno-Blot PVDF membranes. After blocking with 5% bovine serum albumin (BSA) for 1 h at room temperature, the membranes were incubated overnight at 4C with the following primary antibodies: anti-TMEM100 (1:500), anti-TNF-α (1:500), anti-IL-6 (1:500), anti-GNAI2 (1:500), anti-PRDX1 (1:2000). Following three washes with TBST (Tris-buffered saline with 0.1% Tween 20), the membranes were incubated with horseradish peroxidase (HRP)-conjugated secondary antibodies (1:10,000) for 1 h at room temperature. Protein bands were visualized using an enhanced chemiluminescence (ECL) substrate and detected with a Tanon 5200 chemiluminescence imaging system. Band intensities were quantified using ImageJ software, with β-actin serving as the loading control for normalization.

### MTT Colorimetric Assay

Cell proliferation was accessed using an MTT Cell Proliferation Assay Kit. After 24 h of transfection and stimulation, the culture medium was replaced with serum-free medium. Cells were then incubated with MTT solution (5 mg/mL in PBS) for 4 h at 37°C in a 5% CO_2_ atmosphere. Following incubation, the supernatant was carefully removed, and the resulting formazan crystals were dissolved in DMSO with shaking at room temperature for 10 min. Absorbance was measured at 490 nm using a microplate Reader (PerkinElmer, EnSpire).

### EdU Staining Assay

Cell proliferation was evaluated using the BeyoClick™ EdU-555 Cell Proliferation Kit. Following 24 h of transfection and stimulation, cells were incubated with 10 μM EdU for 2 h at 37C under 5% CO_2_. Subsequently, cells were fixed with 4% paraformaldehyde for 15min at room temperature and permeabilized with 0.3% Triton X-100 for 10-15 min, followed by three PBS washes. The click reaction was performed by incubating the cells with a reaction mixture (containing click reaction buffer, CuSO_4_, Azide 555, and click additive solution) for 30 min at room temperature in the dark. Cell nuclei were counterstained with Hoechst 33342, and fluorescent images were acquired using a Zeiss Axio Observer 3 fluorescence microscope.

### Cell Apoptosis Analysis

Cell apoptosis was assessed using the FITC Annexin V Apoptosis Detection Kit I. Following 24 h of transfection and stimulation, cells were detached with 0.25% trypsin (without EDTA) and collected by centrifugation at 1000 rpm for 5 min. After two washes with cold PBS, the cells were resuspended in 1× binding buffer at a density of 1 × 10^6^ cells/mL. A 100 µL aliquot of cell suspension (containing 1 × 10^5^ cells) was transferred to a flow cytometry tube and stained with 5 µL FITC Annexin V and 5 µL propidium iodide (PI). The mixture was gently vortexed and incubated for 15 min at room temperature in the dark. Subsequently, 400 µL of 1× binding buffer was added to each tube, and samples were analyzed with 1 h using a Beckman CytoFLEX flow cytometer.

### Immunofluorescence Staining

Following 24 h of transfection and stimulation, cells grown on sterile coverslips were sequentially fixed with cold methanol for 2 min and 70% ethanol for 5 min. After two washes with PBS, cells were blocked with 1% BSA for 30 min at room temperature, and then incubated with primary antibody (anti-FLAG/HA, 1:100 dilution) for 2 h. Subsequently, cells were incubated with TRITC/FITC-conjugated second antibody (goat anti-mouse/rabbit IgG, 1:100 dilution) for 1 h at room temperature protected from light. After three additional PBS washes, nuclei were counterstained with DAPI (10 µg/ml) for 10 min. Finally, coverslips were mounted onto glass slides using anti-fade fluorescent mounting medium, and images were acquired using a Zeiss LSM980 laser scanning confocal microscope.

### Co-immunoprecipitation Assay

For the immunoprecipitation of endogenous proteins, cells were lysed in beyotime^®^ IP lysis/wash buffer supplemented with protease inhibitors (e.g., 1 mM PMSF). Antibodies were diluted to a concentration of 10 μg/mL in the same IP Lysis/Wash Buffer, and then incubated with protein A/G magnetic beads for 2 h at room temperature with gentle rotation. After magnetic separation, the antibody-coupled beads were incubated with the cell lysates overnight at 4°C with rotation. The beads were subsequently washed three times with IP Lysis Buffer, resuspended in SDS-PAGE loading buffer, and boiled for 5 min before proceeding to western blot analysis.

For the analysis of exogenous proteins, cells were transfected with appropriate plasmids prior to lysis. All subsequent steps were performed identically to the procedure described for endogenous protein immunoprecipitation.

## Statistical Analysis

Data are presented as mean ± standard deviation (SD). Differences between two groups were assessed using an unpaired Student’s t-test. All statistical analyses were performed with GraphPad Prism 10 (GraphPad Software, San Diego, CA, USA). A p-value < 0.05 was considered statistically significant.

## Results

### TMEM100 expression is downregulated in LPS-induced ALI

TMEM100 is highly expressed in normal lung tissues, but is significantly downregulated in the context of lung injury associated with carcinogenesis. To examine the role of TMEM100 in ALI, we first analyzed relevant datasets from the ALI-related gene microarray (GSE2411) available in the GEO database. Applying stringent screening criteria (adjusted *p*-value ≤ 0.001 and logC fold change ≥ 3), we visualized significantly differentially expressed genes (DEGs) by comparing lung tissues from LPS+MV-induced ALI mice with those from control mice.

Venn diagram and volcano plot analyses identified 44 significant differentially expressed genes out of 22,646 analyzed—43 upregulated genes and one downregulated gene, TMEM100 (**Fig. 1A-C**). Cluster analysis and bar chart results further confirmed that TMEM100 expression was significantly lower in lung tissues of ALI compared to controls (*P* < 0.05; **Fig. 1D, E**). We then established an LPS-induced ALI mouse model for pathological assessment. H&E and IHC staining of lung tissues revealed characteristic ALI pathology, including alveolar infiltration of neutrophils and erythrocytes, hyaline membrane formation, and inflammatory cell infiltration **(Fig. 1F)**. Notably, TMEM100 expression was markedly reduced in ALI mice compared to controls (**Fig. 1G**). These findings were further corroborated at both the RNA (**Fig. S1A**) and protein levels (**Fig. S1B**).

**Fig. 1.**
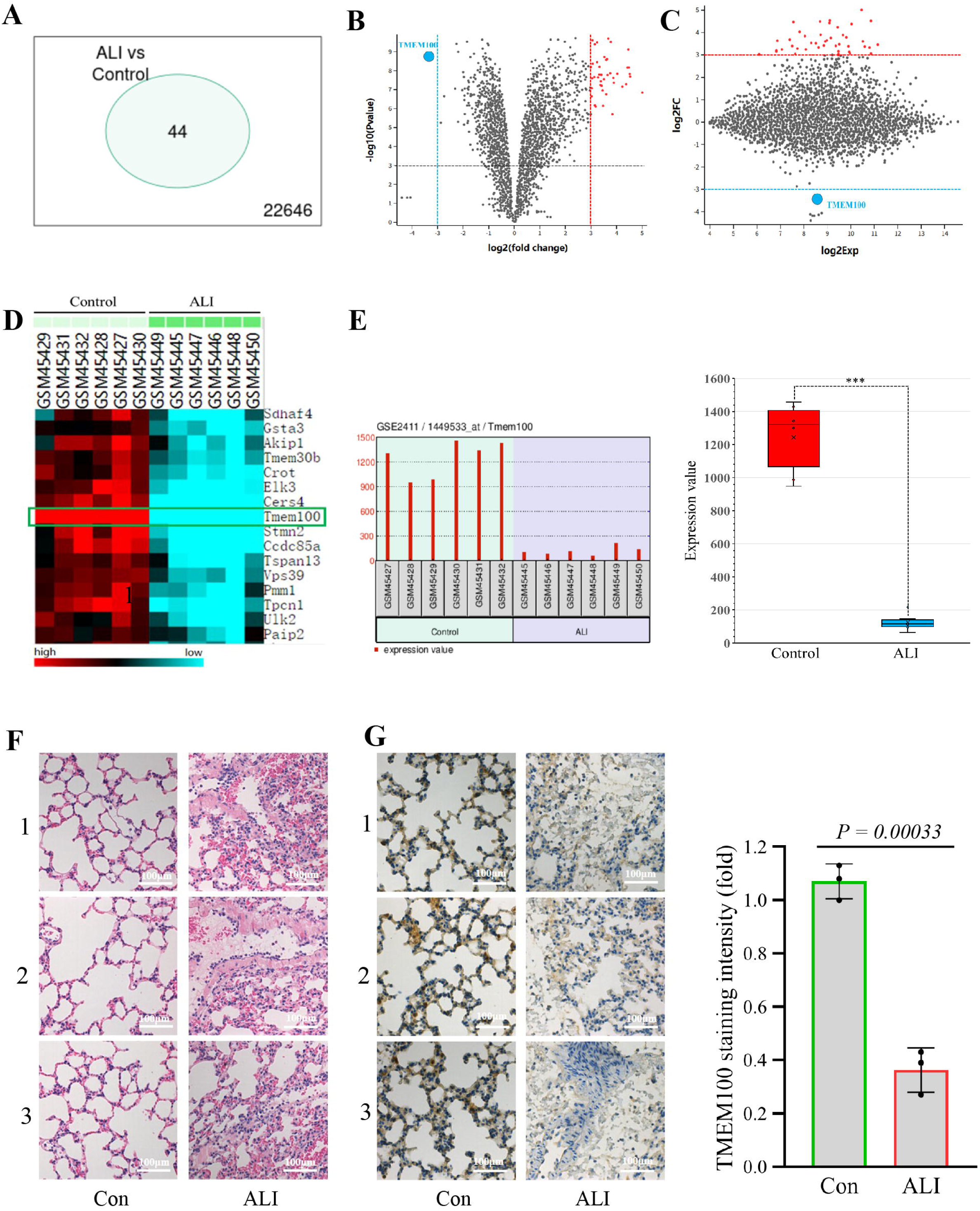
TMEM100 expression is decreased in lung tissues of LPS-induced ALI. **(A)** Venn diagram of differentially expressed genes (DEGs) in lung tissues from mice with LPS+MV-induced ALI and control mice (screening criteria: adjusted *P* ≤ 0.001, |logCFC| ≥ 3). **(B, C)** Volcano plot and mean-difference (MA) plot analysis of DEGs, with blue and red dots indicating downregulated and upregulated genes, respectively. **(D, E)** Cluster heatmap and bar chart of DEGs in individual mice from LPS+MV-induced ALI and control groups (distance metric: uncentered correlations; algorithm: single linkage). **(F)** Representative H&E-stained lung tissue sections from LPS-induced ALI and control mice, showing pathological alterations (scale bar = 100 μm). **(G)** IHC staining of TMEM100 in lung tissues, with brown signal indicating positive expression. Data are presented as mean ± SD; Significance was determined by Student’s t-test: \*\*\**P* < 0.001.

In summary, our results demonstrate significant downregulation of TMEM100 expression in ALI lung tissues, suggesting its potential involvement in the pathogenesis and progression of ALI.

### TMEM100 attenuates LPS-induced lung injury and inflammation in mice

Previous studies have shown that TMEM100 is predominantly expressed in pulmonary endothelial cells and is downregulated in ALI. To investigate its functional role in LPS-induced ALI, we delivered AAV9-TMEM100 or control AAV9-NC to mice via intratracheal instillation. Three weeks after injection, the mice were challenged with LPS, and lung injury was assessed 24 h later. Overexpression of TMEM100 in lung tissue was confirmed by RT-qPCR and western blot analysis (**Fig. S1C, D**).

We first collected lung tissues from each experimental group to evaluate the lung-to-body weight ratio (lung index) and wet/dry weight ratio (W/D ratio). The results showed that LPS stimulation significantly increased both parameters, whereas TMEM100 overexpression substantially attenuated these increases (**Fig. 2A**, **B**). Similarly, LPS challenge elevated total protein levels and total cell counts in bronchoalveolar lavage fluid (BALF), and these effects were significantly suppressed by TMEM100 overexpression (**Fig. 2C**, **D**).

**Fig. 2.**
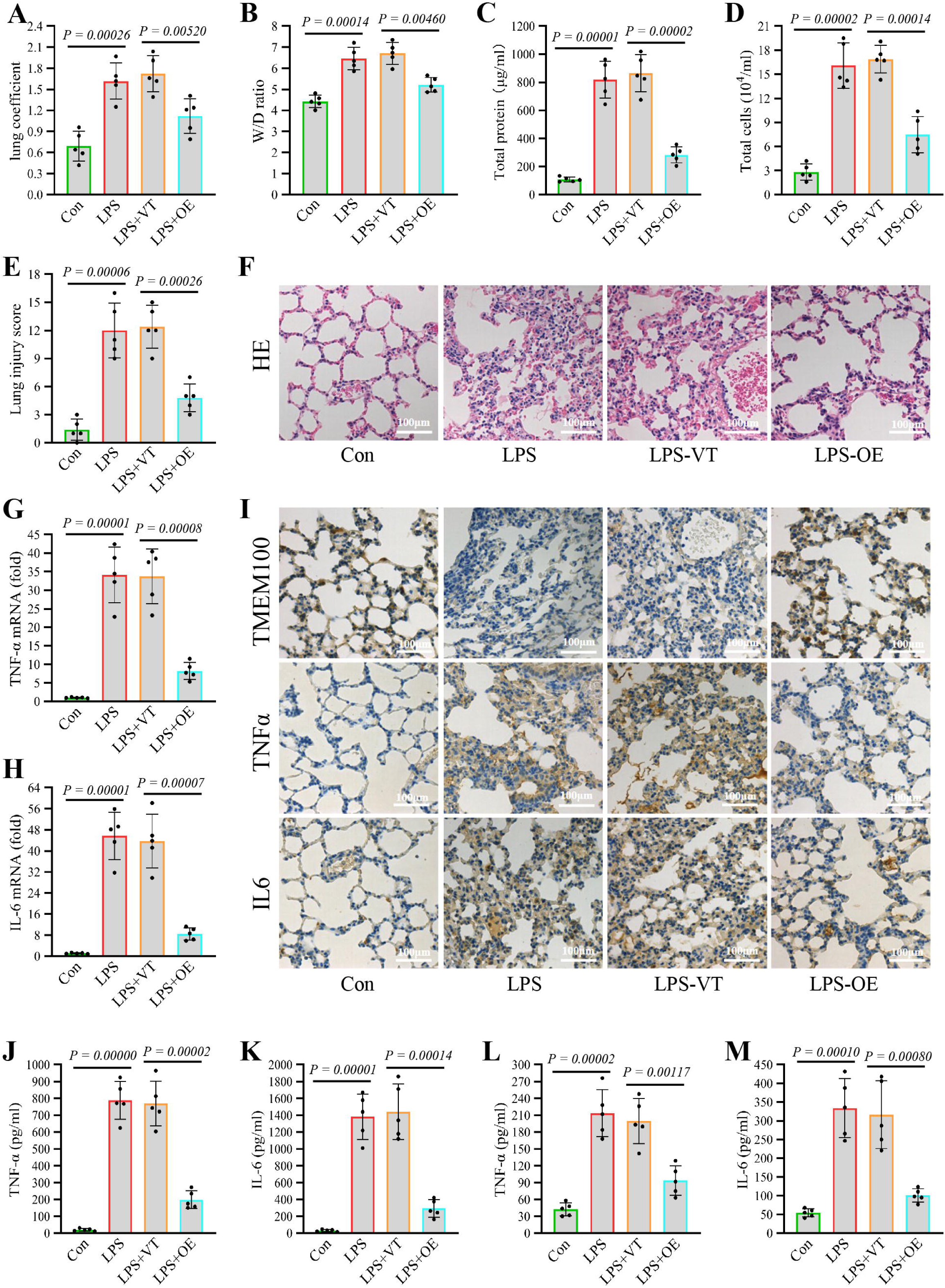
TMEM100 attenuates LPS-induced lung injury and inflammation in mice **(A, B)** Effects of TMEM100 overexpression on the lung index and wet/dry (W/D) ratio in mice with LPS-induced ALI. VT, vector; OE, overexpression. **(C, D)** Effects of TMEM100 overexpression on total protein levels and total cell counts in bronchoalveolar lavage fluid (BALF) from LPS-induced ALI mice. **(E, F)** Effects of TMEM100 overexpression on H&E staining and pathological scores in LPS-induced ALI mice. **(G-I)** RT-qPCR and IHC analysis of TNF-α and IL-6 expression in lung tissues of LPS-induced ALI mice following TMEM100 overexpression. **(J, K)** ELISA measurement of TNF-α and IL-6 levels in BALF and serum from LPS-induced ALI mice with TMEM100 ectopic expression. **(L, M)** ELISA quantification of TNF-α and IL-6 levels in serum from LPS-induced ALI mice with TMEM100 overexpression. All data are presented as mean ± SD; Significance was determined by Student’s t-test.

Histopathological evaluation using H&E staining revealed that LPS induced severe lung injury, characterized by interstitial edema, alveolar wall thickening, and inflammatory cell infiltration. These pathological alterations were markedly ameliorated by TMEM100 overexpression, as further supported by quantitative injury scoring (**Fig. 2E, F**). At the molecular level, RT-qPCR analysis showed that LPS significantly upregulated the mRNA expression of pro-inflammatory cytokines in lung tissues, and these increases were potently inhibited by TMEM100 overexpression (**Fig. 2G, H**). These results were corroborated at the protein level by immunohistochemical staining (**Fig. 2I**).

Consistent with these observations, ELISA results indicated that TMEM100 overexpression significantly reduced the LPS-induced secretion of TNF-α and IL-6 in both BALF and serum (**Fig. 2J-M**). Taken together, these findings demonstrate that TMEM100 overexpression effectively protects against LPS-induced lung injury and inflammation in mice.

### TMEM100 expression is decreased in LPS-induced PVECs

PVECs play a critical role in the pathogenesis of LPS-induced ALI. To assess TMEM100 expression across different lung cell types, we analyzed single-cell RNA sequencing data from the Human Protein Atlas. UMAP visualization and heatmap analysis revealed that TMEM100 expression was not broadly distributed but predominantly localized in endothelial cells (**Fig. 3A, B**), suggesting that endothelial cells may represent the primary target through which TMEM100 regulates ALI.

**Fig. 3.**
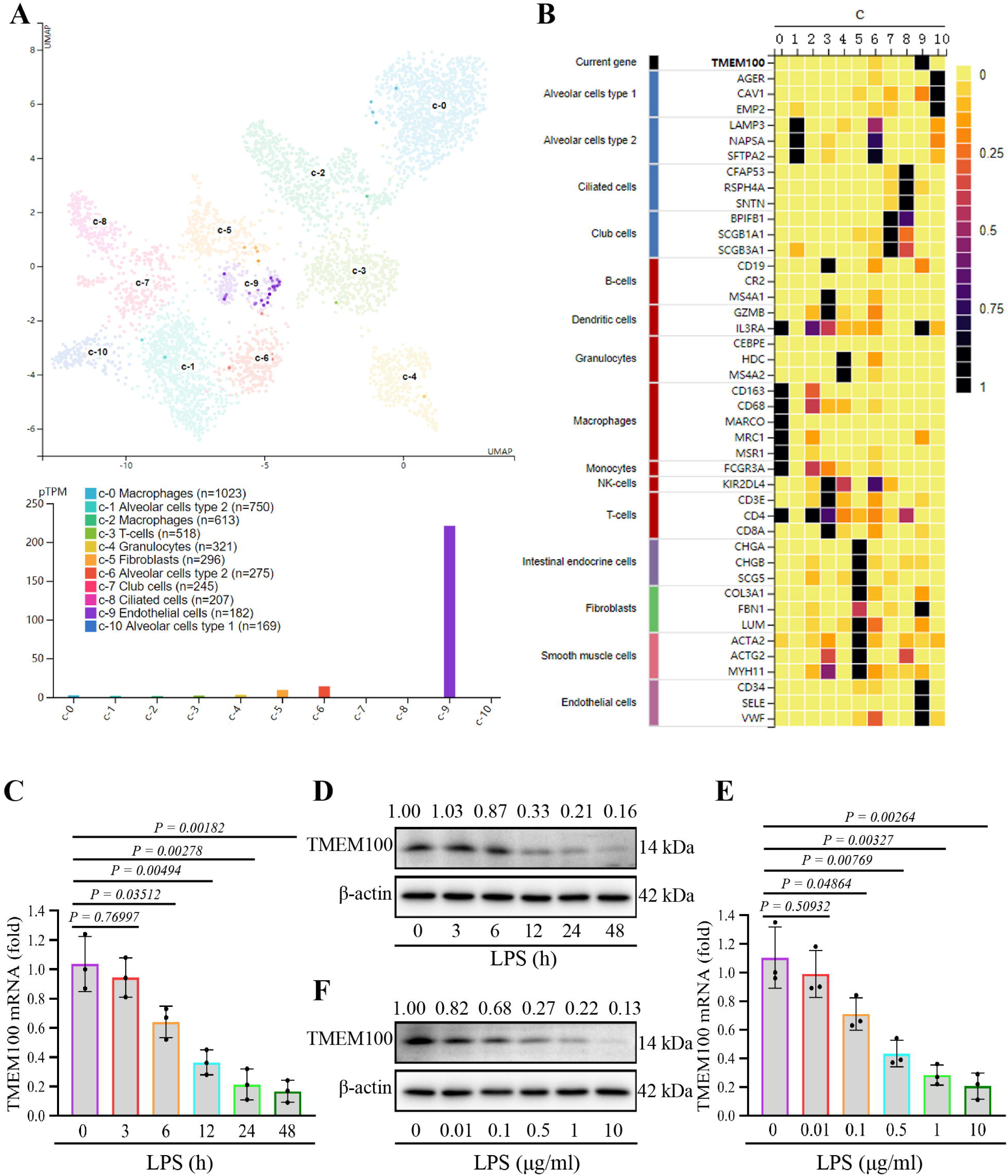
TMEM100 expression is decreased in LPS-induced PVECs **(A)** UMAP visualization of single-cell clusters in lung tissue, with each point representing an individual cell. TMEM100 expression is highlighted across clusters. **(B)** Heatmap analysis of TMEM100 expression levels (top) alongside canonical cell-type markers (left). Color bars indicate functionally related cell types. **(C, D)** RT-qPCR and western blot analysis of TMEM100 mRNA and protein expression in PVECs following LPS stimulation at indicate time points. **(E, F)** RT-qPCR and western blot analysis of TMEM100 mRNA and protein expression in PVECs treated with increasing concentrations of LPS. All data are presented as mean ± SD; Significance was determined by Student’s t-test.

To further explore this, we isolated primary mouse PVECs and stimulated them with LPS to establish an in vitro model of ALI. TMEM100 expression was then evaluated at various time points and LPS concentrations. Our results showed that TMEM100 expression decreased in a time-dependent manner, with the most significant reduction observed 12 h after LPS stimulation (**Fig. 3C, D**). In addition, TMEM100 expression was progressively downregulated with increasing LPS concentrations, reaching the lowest levels at 12 h post-stimulation (**Fig. 3E, F**). These findings demonstrate that TMEM100 expression is downregulated in LPS-induced PVECs in a time-and concentration-dependent manner, implying that TMEM100 may participate in LPS-induced ALI by modulating PVEC activation.

### TMEM100 attenuates the LPS-induced inflammatory response in PVECs

Inflammatory cytokines are key mediators of the inflammatory response. LPS-stimulated PVECs secrete substantial amounts of pro-inflammatory factors and chemokines. To investigate whether TMEM100 overexpression ameliorates LPS-induced inflammation in PVECs, we transfected cells with TMEM100-Flag to transiently elevate its expression (**Fig. 4A**).

**Fig. 4.**
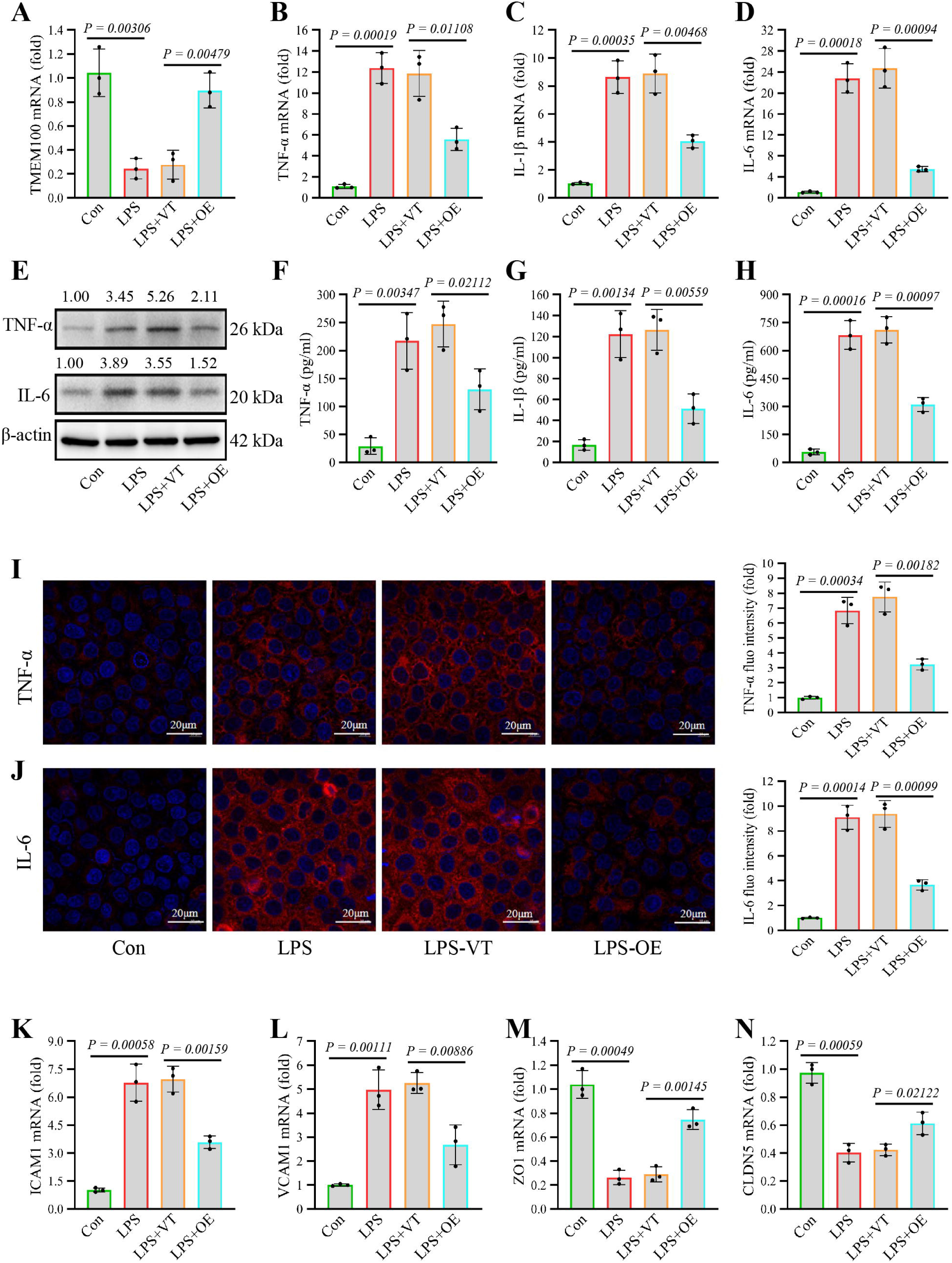
TMEM100 attenuates LPS-induced endothelial cell injury **(A)** RT-qPCR analysis of TMEM100 mRNA expression in PVECs transfected with pcDNA3.1-TMEM100 or emptor vector (pcDNA3.1). **(B-E)** Effect of TMEM100 overexpression on TNF-α, IL-1β and IL-6 expression in LPS-stimulated PVECs, as assessed by RT-qPCR (B-D) and western blot (E). **(F-J)** TMEM100 overexpression reduces the secretion of TNF-α (F), IL-1β (G) and IL-6 (H) by ELISA and decreases their protein levels as detected by immunofluorescence (I, J) in LPS-treated PVECs. **(K-N)** RT-qPCR analysis of ICAM1 (K), VCAM1 (L), ZO1 (M) and CLDN5 (N) expression in LPS-induced PVECs following TMEM100 overexpression. Data are presented as mean ± SD; Significance was determined by Student’s t-test.

RT-qPCR analysis revealed that LPS stimulation significantly upregulated TNF-α, IL-1β and IL-6 mRNA levels in PVECs, which were markedly suppressed by TMEM100 overexpression (**Fig. 4B-D**). Consistent with these findings, western blot and ELISA assays confirmed that TMEM100 overexpression significantly reduced both the expression and secretion of LPS-induced pro-inflammatory cytokines (**Fig. 4E-H**). Immunofluorescence staining further corroborated these findings (**Fig. 4I, J**).

Cell adhesion molecules, such as ICAM1 and VCAM1, play a critical role in neutrophil adhesion to activated endothelium. RT-qPCR demonstrated that TMEM100 overexpression significantly attenuated LPS-induced ICAM1 and VCAM1 expression (**Fig. 4K, L**). Moreover, tight junction proteins (ZO1 and Claudin5) are essential for maintaining PVEC barrier integrity, and their dysregulation is closely associated with endothelial injury. Notably, TMEM100 overexpression restored the expression of ZO1 and Claudin5 following LPS stimulation (**Fig. 4M, N**). In summary, these findings demonstrate that TMEM100 exerts a protective effect against LPS-induced inflammatory activation and monocyte adhesion in PVECs.

### TMEM100 mitigates LPS-induced impairment of proliferation and apoptosis in PVECs

Under physiological conditions, PVECs maintain vascular homeostasis through a tightly regulated balance between proliferation and apoptosis. However, LPS disrupts this equilibrium by suppressing proliferation and promoting apoptosis. EdU staining showed that LPS stimulation significantly inhibited PVEC proliferation, and this suppression was substantially attenuated by TMEM100 overexpression (**Fig. 5A**). This finding was corroborated by MTT assay (**Fig. 5B**), which showed that TMEM100 overexpression effectively counteracted the LPS-induced proliferation inhibition.

**Fig. 5.**
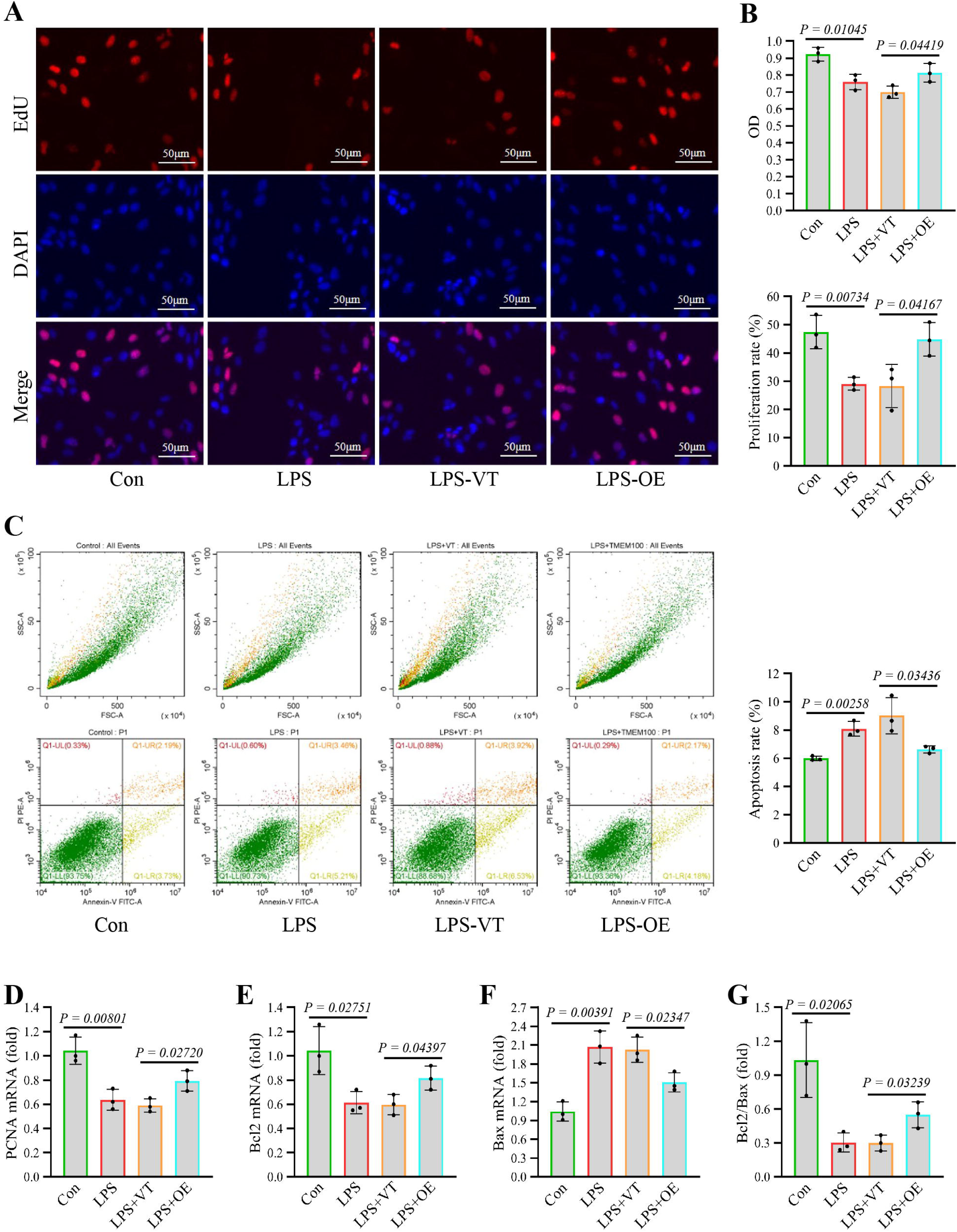
TMEM100 restores the proliferation-apoptosis balance in PVECs **(A)** EdU assay showing proliferating cells (red fluorescence) in LPS-induced PVECs with or without TMEM100 overexpression. **(B)** Cell proliferation assessed by MTT assay in LPS-induced PVECs following TMEM100 ectopic expression. **(C)** Apoptosis analysis by flow cytometry in LPS-induced PVECs. The total apoptosis rate represents the combined proportion of early (Q1-LR) and late (Q1-UR) apoptotic cell populations. **(D-G)** RT-qPCR analysis of PCNA (D), Bcl-2 (E), Bax (F) expression and Bcl-2/Bax ratio (G) in LPS-induced PVECs upon TMEM100 overexpression. Data are presented as mean ± SD; Significance was determined by Student’s t-test.

Flow cytometry analysis revealed that LPS stimulation markedly increased PVEC apoptosis, as expected, while TMEM100 overexpression significantly attenuated this pro-apoptotic response (**Fig. 5C**). Furthermore, RT-qPCR analysis of proliferation-and apoptosis-related markers (PCNA, Bcl-2, and Bax) confirmed these observations at the mRNA level (**Fig. 5D-G**). Collectively, these results demonstrate that TMEM100 effectively restores the balance between proliferation and apoptosis in LPS-treated PVECs.

### TMEM100 attenuates LPS-induced NF-**κ**B activation in PVECs

The NF-κB signaling pathway plays a critical role in regulating inflammatory response, cell proliferation and apoptosis. Its aberrant activation is a key mechanism underlying LPS-induced ALI. Western blot analysis revealed that LPS significantly increased the phosphorylation of IκBα and p65 in PVECs, which was markedly suppressed by TMEM100 ectopic expression (**Fig. 6A**).

**Fig. 6.**
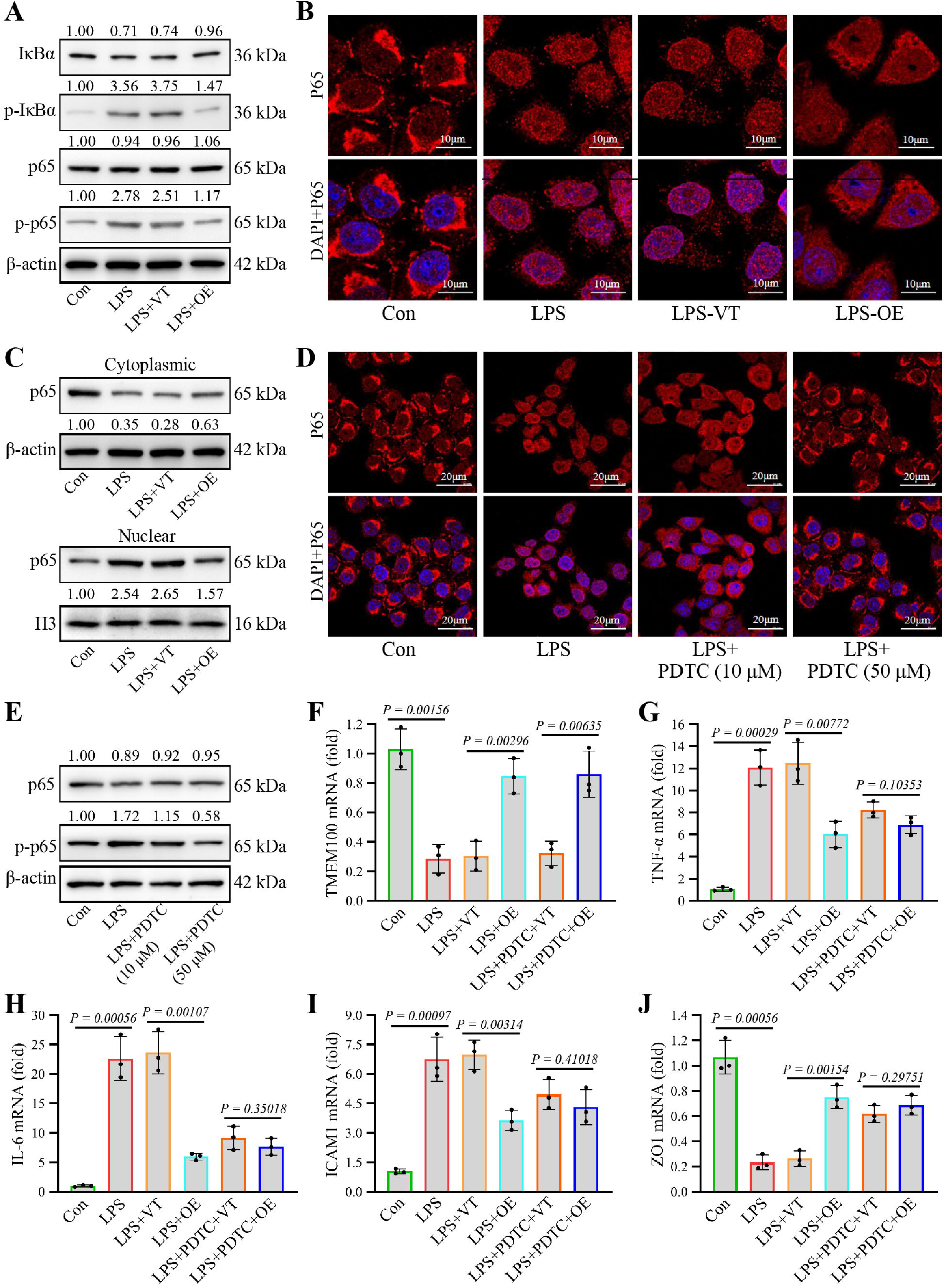
TMEM100 overexpression suppresses LPS-induced NF-**κ**B activation **(A)** Western blot analysis of IκBα and p65 phosphorylation in LPS-stimulated PVECs following TMEM100 overexpression. **(B)** Immunofluorescence staining of p65 nuclear translocation in LPS-stimulated PVECs with or without TMEM100 ectopic expression. **(C)** Western blot analysis of p65 expression in cytoplasmic and nuclear fractions from LPS-treated PVECs upon TMEM100 overexpression. **(D, E)** Immunofluorescence staining and western blot analysis of p65 nuclear translocation and phosphorylation levels in LPS-treated PVECs with NF-κB inhibitor (PDTC) treatment. **(F-J)** RT-qPCR analysis of TMEM100 (F), TNF-α (G), IL-6 (H), ICAM1 (I) and ZO1 (J) mRNA expression in LPS-treated PVECs upon TMEM100 overexpression and/or PDTC treatment. Data are presented as mean ± SD; Significance was determined by Student’s t-test.

To assess nuclear translocation of p65, we performed subcellular fractionation followed by immunofluorescence staining, which showed that LPS stimulation reduced cytoplasmic p65 levels while increasing nuclear accumulation of p65. Notably, TMEM100 overexpression blocked LPS-induced nuclear translocation of p65 in PVECs (**Fig. 6B**). This finding was further confirmed by western blot analysis of nuclear and cytoplasmic fractions (**Fig. 6C**), collectively demonstrating that TMEM100 inhibits LPS-induced NF-κB activation.

To determine whether anti-inflammatory effects of TMEM100 depend on NF-κB signaling, we used pyrrolidinedithiocarbamic (PDTC), a specific NF-κB inhibitor, to block pathway activation (**Fig. 6D, E**). Subsequent analysis showed that the ability of TMEM100 to suppress the expression of inflammatory cytokines, cell adhesion molecules and tight junction proteins was significantly attenuated or abolished when NF-κB signaling was inhibited (**Fig. 6F-J**). Further investigation showed that Diprovocim, an NF-κB signaling pathway activator, reversed the suppression of inflammatory factors (TNF-α and IL-6) induced by TMEM100 overexpression (**Fig. S2**). These results indicate that TMEM100 exerts its protective effects in ALI primarily through modulation of the NF-κB signaling pathway.

### TMEM100 disrupts the PRDX1/GNAI2 complex

To elucidate the molecular mechanism through which TMEM100 regulates NF-κB signaling, we conducted a series of protein-protein interaction assays. We first performed co-immunoprecipitation (Co-IP) using an anti-Flag antibody to isolate TMEM100-associated proteins from PVECs expressing Flag-TMEM100. Subsequent mass spectrometry analysis identified several candidate binding partners, including PRDX1 and GNAI2—both known regulators of NF-κB signaling (**Fig. 7A, B**). Molecular docking simulations further supported the potential binding of TMEM100 to PRDX1 and GNAI2 (**Fig. 7C, D**).

**Fig. 7.**
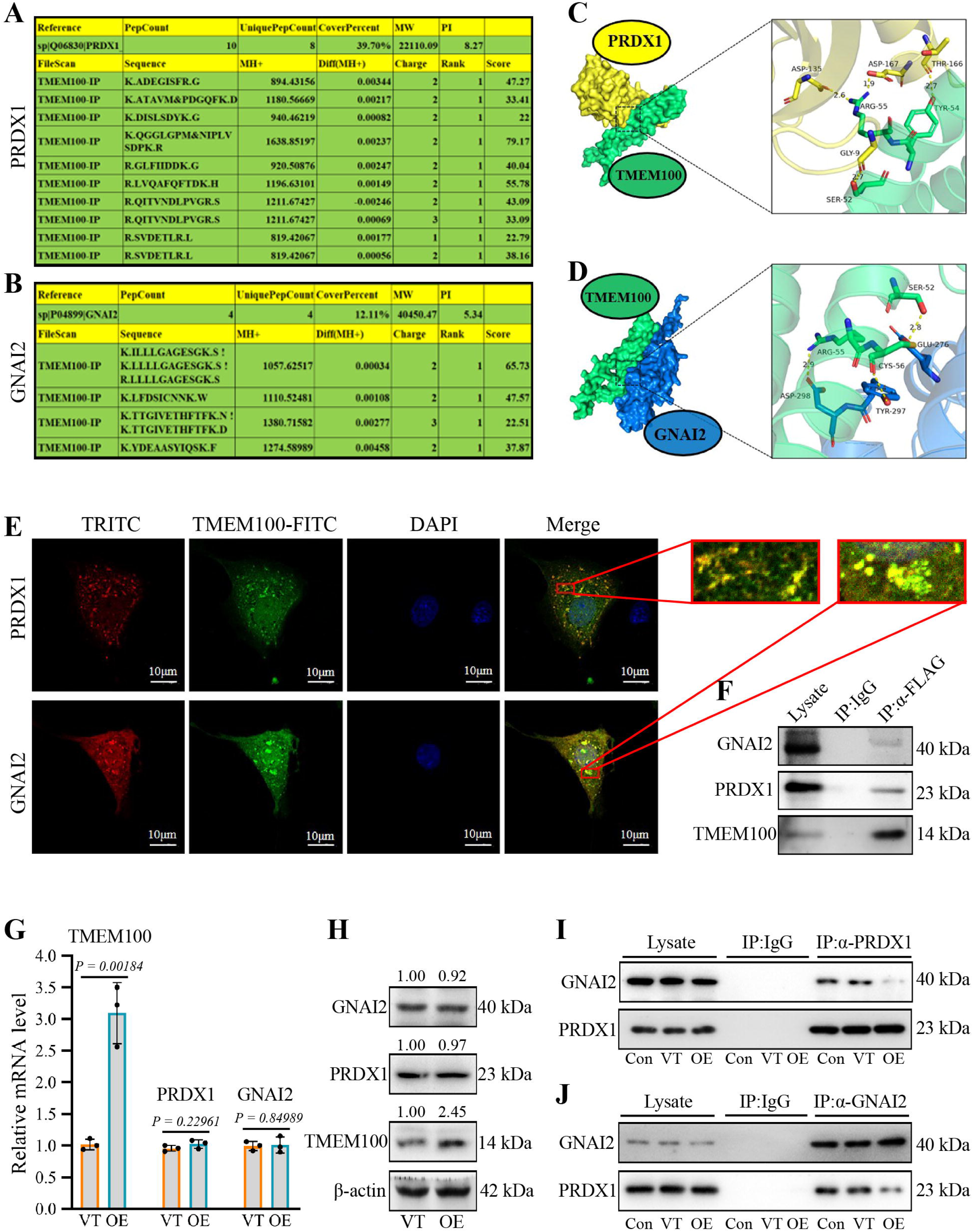
TMEM100 disrupts the PRDX1/GNAI2 complex **(A, B)** Mass spectrometry analysis of anti-Flag immunoprecipitates identifing PRDX1 and GNAI2 as potential TMEM100-interacting proteins. **(C, D)** Molecular docking models showing potential binding conformations between TMEM100 and PRDX1 or GNAI2. **(E)** Immunofluorescence images showing cytoplasmic co-localization of TMEM100 with PRDX1 and GNAI2 in PVECs. **(F)** Co-IP assays confirming TMEM100 interaction with PRDX1/GNAI2 in vivo (using anti-TMEM100 for IP, IgG as negative control). **(G, H)** RT-qPCR analysis and western blot analysis of PRDX1 or GNAI2 upon TMEM100 overexpression. **(I, J)** Co-IP assays between PRDX1-GNAI2 upon TMEM100 overexpression (using anti-PRDX1 or GNAI2 for IP, IgG as negative control). Data are presented as mean ± SD; Significance was determined by Student’s t-test.

To validate these interactions, we co-transfected Flag-TMEM100 with HA-PRDX1 or HA-GNAI2 into PVECs and examined their subcellular localization. Immunofluorescence staining revealed strong co-localization of TMEM100 with both PRDX1 and GNAI2 (**Fig. 7E**), suggesting possible physical interaction. Direct binding was further confirmed by Co-IP assays (**Fig. 7F**).

Notably, RT-qPCR analysis showed that TMEM100 overexpression did not alter PRDX1 or GNAI2 mRNA levels (**Fig. 7G, H**), suggesting that TMEM100 acts at the post-translational level. To determine whether TMEM100 affects the stability of PRDX1-GNAI2 complex, we performed Co-IP using a PRDX1-specific antibody. We observed that although PRDX1 and GNAI2 readily co-immunoprecipitated under basal conditions, TMEM100 overexpression markedly reduced their binding affinity (**Fig. 7I, J**). These findings suggest that TMEM100 competitively binds to PRDX1, destabilizing the PRDX1-GNAI2 complex and consequently suppressing NF-κB activation. This mechanism provides a molecular basis for the TMEM100’s anti-inflammatory effects in LPS-induced ALI.

## Discussion

ALI is a severe inflammatory pulmonary condition characterized by increased alveolar-capillary barrier permeability, resulting from diverse intra-and extra-pulmonary (non-cardiogenic) pathogenic factors. LPS, the bioactive component of endotoxin, represents a major etiological factor in ALI development. The pathogenesis of LPS-induced ALI involves multiple interrelated mechanisms, with inflammation being the most extensively studied, along with oxidative stress and apoptosis. The inflammatory cascade initiates when LPS binds to lipopolysaccharide-binding protein (LBP), forming an LPS-LBP complex that subsequently associates with CD14. This ternary complex triggers robust activation of inflammatory signaling pathways in immune cells, resulting in excessive release of pro-inflammatory cytokines and consequent cytokine dysregulation. Notably, PVECs serve as primary targets of LPS-mediated injury. Their dysfunction constitutes a fundamental pathological feature underlying ALI initiation and progression [22]. Therefore, targeting the dysfunction of endothelial cells is an important strategy for ALI [23–25].

In this study, we initially analyzed gene expression profiles in an ALI mouse model and identified multiple differentially expressed genes, including TMEM100. TMEM100 is known to play a significant role in lung cancer progression and exhibits potential anti-tumor activity. Database analyses further indicate TMEM100 contributes to lung development and functional maintenance. We observed a significant decrease in TMEM100 expression in lung tissues from LPS-induced ALI mice. Furthermore, we found that TMEM100 is specifically expressed in PVECs, and its expression was markedly downregulated following LPS stimulation in both concentration-and time-dependent manners.

TNF-α is aberrantly secreted by PVECs during ALI, and triggers the release of downstream inflammatory factors, such as IL-1β, IL-6, and IL-8, thereby amplifying the inflammatory cascade [26]. IL-6 exacerbates ALI by directly damaging endothelial cells and increasing vascular permeability. Activated neutrophils contribute to alveolar dysfunction and aggravate lung injury through mediator release, degranulation, and chemotaxis [27]. Our findings demonstrated that TMEM100 effectively attenuated inflammatory cascades, neutrophil infiltration, and lung tissue damage in ALI, suggesting its potential as a therapeutic target for mitigating disease progression.

Under physiological conditions, PVECs maintain vascular homeostasis through a tightly regulated balance between proliferation and apoptosis. However, LPS disrupts this equilibrium by suppressing proliferation and promoting apoptosis, leading to increased vascular permeability, neutrophil infiltration into pulmonary stroma and alveoli, and ultimately respiratory dysfunction. These pathological changes exacerbate lung injury and elevate mortality in ALI. Our findings demonstrate that TMEM100 effectively counteracts LPS-induced dysregulation of PVEC proliferation and apoptosis, thereby preserving endothelial integrity.

The NF-κB signaling pathway plays a central role in modulating inflammation, cell proliferation, and apoptosis. Its aberrant activation is a key driver of LPS-induced ALI. Our preliminary mass spectrometry data suggested that TMEM100 may interact with GNAI2 and/or PRDX1. Studies have demonstrated that GNAI2 serves as a key mediator of inflammation in liver ischemia-reperfusion injury, alveolar and airway allergic inflammation, and LPS-induced lung injury [28]. As a molecular chaperone, PRDX1 modulates intracellular transcription levels and oxygen-free radical formation, playing a significant role in inflammation regulation [29]. Notably, PRDX1 binds to TRAF6 (TNF receptor-associated factor 6), inhibiting its ubiquitin ligase activity and reducing the ubiquitination of ECSIT (evolutionarily conserved signaling intermediate in Toll pathway), a critical step in NF-κB pathway activation [30]. Additionally, evidence suggests that GNAI2 competitively binds PRDX1, displacing TRAF6 and enhancing its ubiquitin ligase activity. This increases ECSIT ubiquitination, thereby activating the TLR4 (Toll-like receptor 4)-mediated NF-κB pathway and amplifying inflammatory responses [31]. Intriguingly, we found that TMEM100 binds to both GNAI2 and PRDX1 simultaneously, yet its overexpression did not affect the expression levels of either protein. Subsequent Co-IP results demonstrate that TMEM100 significantly reduces GNAI2-PRDX1 binding, revealing a potential mechanism by which it suppresses NF-κB pathway activation.

In summary, this study first identified decreased TMEM100 expression in LPS-induced ALI through bioinformatic analysis, a finding subsequently validated in an animal model. We further demonstrated that TMEM100 overexpression exerts protective effects both in vivo and in vitro. Finally, we confirmed that the underlying mechanism involves TMEM100-mediated disruption of the PRDX1-GNAI2 interaction, which in turn regulate of the activation of NF-κB signaling—a central pathway in LPS-induced ALI. These results underscore the protective role of TMEM100 in endotoxin-induced ALI and provide a theoretical foundation for understanding its biological functions and potential relevance to ALI gene therapy.

## Acknowledgements

Thanks for the support of Center for Scientific Research, Anhui Medical University.

---

**Fig. S1 Validation of TMEM100 expression and its effect in mouse lungs**

(**A, B**) TMEM100 mRNA (A) and protein (B) expression levels in lung tissues from control and ALI mice. (**C, D**) Efficiency of TMEM100 overexpression in lung tissues, as assessed by RT-qPCR (C) and Western blot (D) analysis.

**Fig. S2 ELISA analysis of TNF-**α **(A) and IL-6 (B) secretion from LPS-induced PVECs with or without TMEM100 overexpression.**

