## Supplementary material for "TMEM100 attenuates NF-κB activation via disrupting the PRDX1-GNAI2 complex to alleviate acute lung injury": Table S1

**Table S1. Sequences of Oligo-primers used in this study**

| **Number** | **Sequence (5’-3’)** | **Purpose** |
| --- | --- | --- |
| P1 | TGCCGAACTCTCCTGCTACCG | Primers for qRT-PCR of Mouse TMEM100 |
| P2 | GCCGAAGATGGAGATGATGGAACC |  |
| P3 | TTCTCATTCCTGCTTGTGG | Primers for qRT-PCR of Mouse TNF-α |
| P4 | ACTTGGTGGTTTGCTACG |  |
| P5 | CTTCTTGGGACTGATGCTGGTGAC | Primers for qRT-PCR of Mouse IL-6 |
| P6 | AGTGGTATCCTCTGTGAAGTCTCCTC |  |
| P7 | TCACTATTGGCAACGAGCGGTTC | Primers for qRT-PCR of Mus actin |
| P8 | GGCATAGAGGTCTTTACGGATGTCAAC |  |
| P9 | GTACCGAGCTCTCCTGCTACC | Primers for qRT-PCR of Human TMEM100 |
| P10 | CCAGGCCAAAGATGGAGATA |  |
| P11 | CTCCTCACCCACACCATCA | Primers for qRT-PCR of Human TNF-α |
| P12 | GGAAGACCCCTCCCAGATAG |  |
| P13 | TTCGGTCCAGTTGCCTTCT | Primers for qRT-PCR of Human IL-6 |
| P14 | GGTGAGTGGCTGTCTGTGTG |  |
| P15 | TGGCAATGAGGATGACTTGT | Primers for qRT-PCR of Human IL-1β |
| P16 | TGGTGGTCGGAGATTCGTA |  |
| P17 | GGCCGGCCAGCTTATACAC | Primers for qRT-PCR of Human ICAM1 |
| P18 | TAGACACTTGAGCTCGGGCA |  |
| P19 | TCAGATTGGAGACTCAGTCATGT | Primers for qRT-PCR of Human VCAM1 |
| P20 | ACTCCTCACCTTCCCGCTC |  |
| P21 | AACTGGGCTCTTGGCTTGCTATTC | Primers for qRT-PCR of Human ZO1 |
| P22 | TCCAGAAGTCAGCACGGTCTCC |  |
| P23 | CGCCTTCCTGGACCACAACATC | Primers for qRT-PCR of Human CLDN5 |
| P24 | AGAGCCAGCACCGAGTCGTAC |  |
| P25 | CTCTTCCAGCCTTCCTTCCT | Primers for qRT-PCR of Human actin |
| P26 | AGCACTGTGTTGGCGTACAG |  |
