## Supplementary figures and images for "TMEM100 attenuates NF-κB activation via disrupting the PRDX1-GNAI2 complex to alleviate acute lung injury"

### Fig S1

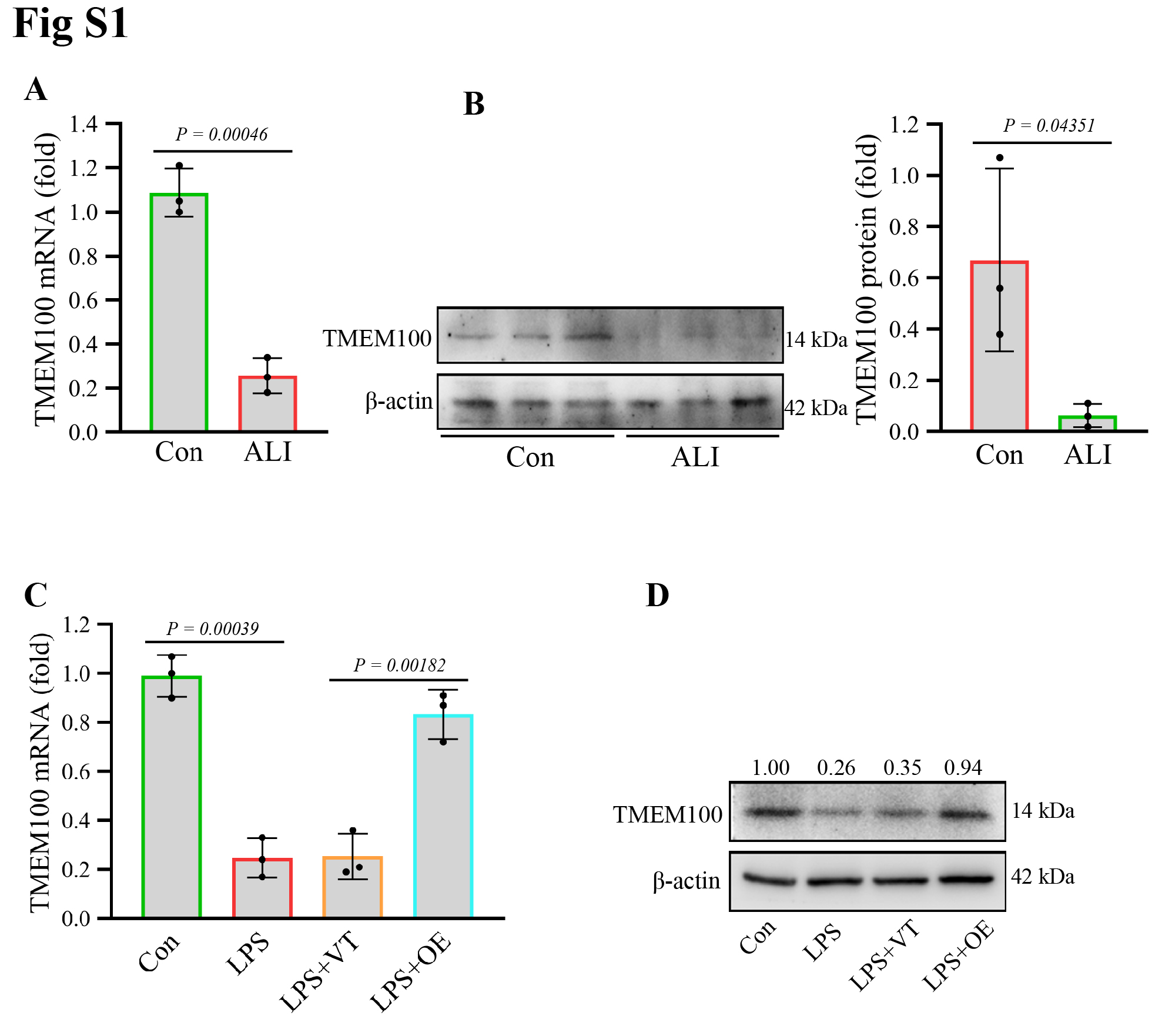

### Fig S2

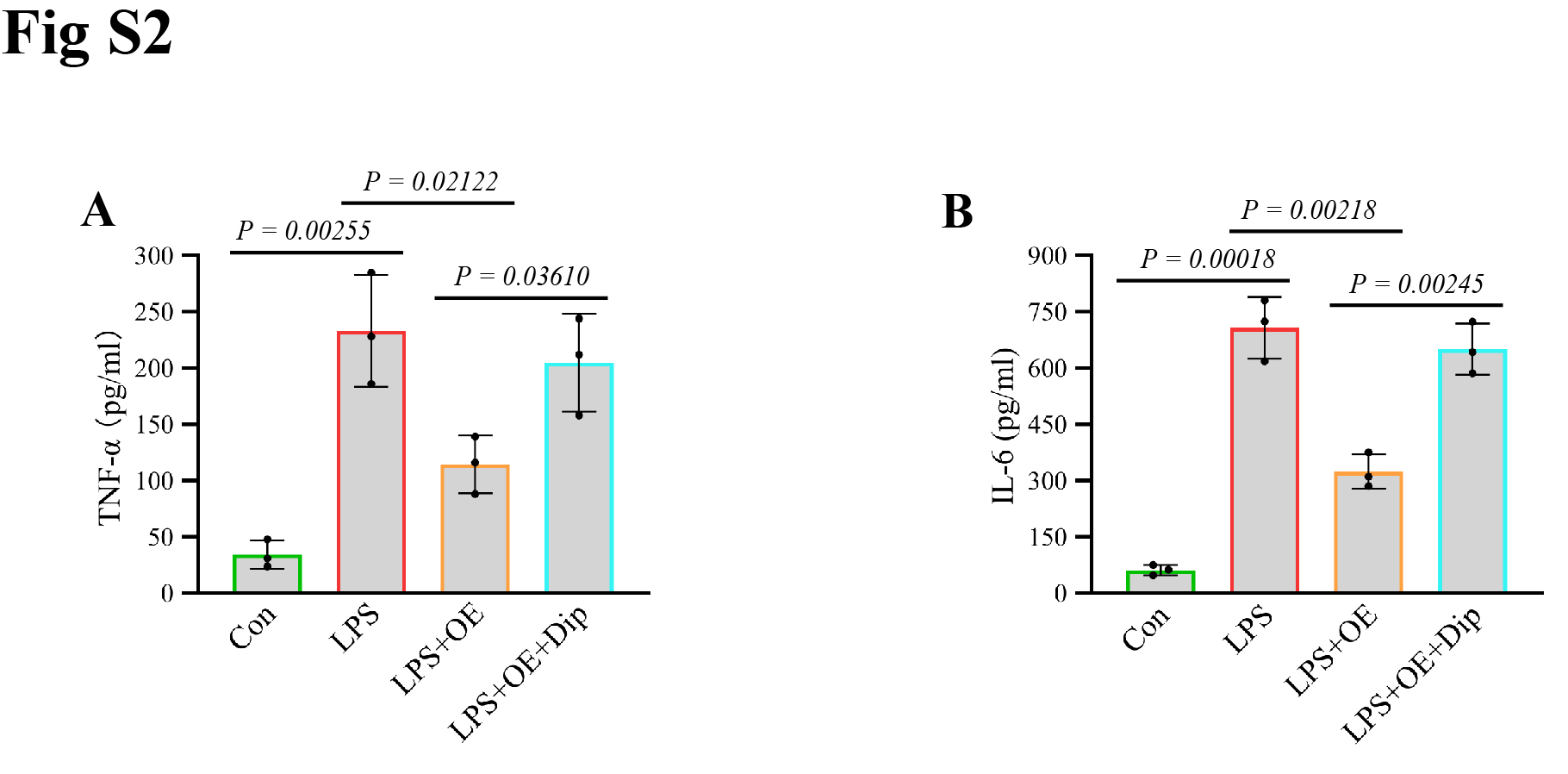
